# Chemotherapy reprograms the breast cancer pericyte secretome from growth-promoting to pro-inflammatory

**DOI:** 10.64898/2026.11.25.769223

**Authors:** Kevin Muñoz Forti, Maggie H. Chassé, Hardik Shah, Natalia Antonides-Jensen, Sonia L. Hernandez, John C. Alverdy, Samantha S. Yee

## Abstract

**Background:** Breast cancer (BrCa) is one of the most common cancers in the United States, and metastatic disease has a 5-year survival rate around 30%. Taxanes such as docetaxel (DTX) are standard of care, but chemoresistance occurs in 30-35% of patients. The tumor microenvironment (TME) shapes both progression and therapeutic response. Pericytes regulate vascular stability and angiogenesis, resemble cancer-associated fibroblasts, and are exposed to chemotherapy in the TME. The metabolites that pericytes secrete, and how chemotherapy changes them, are largely unexplored. We hypothesized that the pericyte secretome promotes BrCa proliferation and that docetaxel shifts it toward a pro-inflammatory profile.

**Methods:** We cultured three BrCa cell lines with pericyte-conditioned media (CM) and measured proliferation and migration. We profiled cancer cells cultured with CM from DTX-treated or vehicle-treated pericytes by RNA sequencing, analyzed CM by polar metabolomics, and modeled recipient-cell metabolism.

**Results:** CM increased proliferation and migration in all three cell lines and induced a high-grade-like program marked by NNMT and SERPINE1/PAI-1. DTX-CM did not globally alter the transcriptome but added a SASP-like TNFα-NFκB program. DTX also raised extracellular uridine, thymidine and cytidine in CM. Undiluted DTX-CM reduced proliferation in all three lines, whereas pericyte CM did not shift the docetaxel IC50. Exploratory metabolic modeling predicted higher extracellular transport and lower nucleotide interconversion in recipient cancer cells.

**Conclusion:** Docetaxel shifts the pericyte secretome from growth-promoting to pro-inflammatory without blunting docetaxel potency in vitro. Pericyte-derived nucleosides and SASP-like cytokines are candidate mediators of this shift.

## INTRODUCTION

Breast cancer (BrCa) is one of the most common cancers for women, and its progression and response to therapy depend on the tumor microenvironment (TME) as well as on tumor cells [1]. Docetaxel (DTX) and other taxanes are the standard of care, yet chemoresistance occurs in 30-35% of patients and metastatic disease has a 5-year survival of account 30% [2, 3]. Recurrence and metastasis therefore persist despite advances in therapy, and there is an unmet need to better understand how non-cancer cells contribute to this process. The TME consists of several cell types and while some have been explored, many remain poorly defined.

Pericytes are mural cells that encase endothelial cells in capillaries and venules and regulate vascular stability and angiogenesis [4]. They resemble cancer-associated fibroblasts and respond dynamically to their environment, including chemotherapy [5, 6]. Cancer-associated fibroblasts have been suggested to arise from cellular sources including pericytes [7, 8]. Pericytes secrete bioactive factors that influence angiogenesis, immune modulation, and tumor cell survival [9, 10]. We previously found that docetaxel enriched interleukin-6 (IL-6), a pro-inflammatory cytokine implicated in pro-survival mechanisms, in the pericyte secretome [11]. This suggested that docetaxel alters pericyte secretory programs. What remains unknown is the full composition of the pericyte secretome, how docetaxel alters it, and how these changes affect cancer cells.

We hypothesized that pericytes support BrCa growth and migration through their secretome, and that docetaxel converts this growth-promoting secretome into a pro-inflammatory state. To test this, we cultured ER+ and triple-negative BrCa cells with conditioned media from vehicle-treated or DTX-treated pericytes. We profiled the transcriptional response of cancer cells to conditioned media, the metabolome of the conditioned media, and the predicted metabolic state of recipient cells. Here we show that pericyte conditioned media supports cancer cell growth and migration, and that docetaxel converts it into media that induces an inflammatory, SASP-like response in cancer cells and is enriched for pyrimidine nucleosides, without altering cancer cell sensitivity to docetaxel in vitro. Overall, these findings suggest that chemotherapy rewires pericyte-tumor communication from growth support to inflammation.

## MATERIALS & METHODS

### Reagents

Docetaxel (DTX) (#01885-5MG-F) was purchased from Sigma Aldrich, and DMSO, which is the vehicle control solvent for DTX, was purchased from ThermoFisher (#J66650.AK). In addition, HyClone™ Water, Molecular Biology Grade (#SH3053802) was bought from Cytiva for RNA elution. 1x DPBS was purchased from ThermoFisher (#14190144). 16% paraformaldehyde was utilized for dilution to 4% Paraformaldehyde (PFA) (ThermoFisher; #PI28908). All other reagents are described in the text below.

### Cell Culture

We obtained MCF-7 (HTB-22, RRID: CVCL_0031) and T-47D (HTB-133, RRID: CVCL_0553) cancer cell lines from ATCC. The MDA-MB-231 (HTB-26, RRID: CVCL_0062) cell line was obtained from Dr. Jinyang Li in the laboratory of Dr. Anthony Kossiakoff (University of Chicago). Human Brain Vascular Pericytes (HBVP) were purchased from ScienCell Research Laboratories (#1200; lot number 32562, previously #1020), maintained in ScienCell Pericyte Medium (ScienCell Research Laboratories #1201), and cultured up to passage 10 based on consistent doubling times and cell morphology and health. Heat-inactivated fetal bovine serum (FBS) was used for cell culture (Life Technologies #A5669701; Thermo Fisher Scientific #1600-044). We used Dulbecco’s Modified Eagle Medium (DMEM) (Thermo Fisher Scientific #10-013-CV) supplemented with 10% FBS and 1% penicillin-streptomycin (P/S; Gibco #15-140-122) to culture all cancer cell lines. The MDA-MB-231 cell line was maintained initially in 1:1 DMEM supplemented with 10% FBS and 1% P/S and Roswell Park Memorial Institute (RPMI) 1640 medium (Thermo Fisher Scientific #11-875-119) with 10% FBS and 1% P/S, and the MCF-7 cell line was maintained in Improved Minimum Essential Medium (IMEM) (Corning #A1048901) with 10% FBS and 2.5 mL of 10 mg/mL gentamicin (Gibco #15710064) in 500 mL. Cells were cultured at 37°C in a humidified incubator with 5% CO₂ and were routinely tested for mycoplasma contamination (Invivogen #rep-mys-50). A brief characterization of the human cells utilized for this study is in Supplementary Table 1 from [11].

### Pericyte-conditioned media without pre-treating the pericytes

When pericytes reached a confluency of 70-90%, cells were washed with 1 x DPBS prior to the addition of serum-free media (0.1% Bovine serum albumin (Millipore Sigma/Roche #3117057001), 1% P/S for 24 hours). Conditioned media was collected, cleared by centrifugation for 5 minutes at 600g at room temperature (RT), and stored at -80 °C or used immediately.

### Pericyte conditioned media with pre-treating the pericytes

To minimize cross-contamination of DTX in the conditioned media, we generated conditioned media as previously described. Briefly, when pericytes reached 70-90% confluency, they were treated with DMSO (vehicle control) or 10 nM DTX for 24 hours in complete growth media. After 24 hours, the media were removed, the wells were washed with PBS, and the cells were refed with serum-free media (0.1% BSA (Millipore Sigma/Roche #3117057001) and 1% P/S) for 24 hours before collection of the conditioned media. The collected conditioned media was then centrifuged at 600g to clear dead cells or cell debris and either used immediately or aliquoted and frozen at -80°C.

### Cell Counting Kit-8 assay

Approximately 1,000 MCF-7, 4,000 MDA-MB-231, or 7,000 T-47D breast cancer cells were seeded per well into a 96-well plate in 100 µL of their respective growth media. The cells were allowed to adhere overnight. The following day, the media were removed and replaced with 1) 200 µL control media (DMEM SFM) or pericyte-conditioned media, 2) 200 µL control media (DMEM SFM) or pericyte-conditioned media with and without exogenous docetaxel or DMSO control, or 3) 200 µL control pericyte-conditioned media or 10 nM DTX pericyte-conditioned media in a 1:1, 1:10, and 1:100 dilution series. After 72 hours of incubation, 20 µL CCK-8 (Dojindo #CK04-11/Thermo Fisher Scientific #NC9864731) was added to each well for 3 – 4 hours at 37°C. The 96-well plates were read at 450 nm using the SpectraMax i3x plate reader (Molecular Devices). Absorbance values at 450 nm were calculated by subtracting the background from media-only wells from the acquired data. GraphPad Prism software (version 10) was used to visualize the data.

### Migration assay

Transwell inserts (Thermo) with 8 µm pores were placed in 24-well plates containing 900 µL of control media or pericyte-conditioned media and preconditioned in a tissue culture incubator for 20 min. Approximately 100,000 live cells (MCF-7, MDA-MB-231, or T-47D cell lines) were seeded into the preconditioned insert in a total of 300 µL DMEM SFM. Each condition was conducted in technical duplicates for each independent experiment. After 24 hours, cells in the top chamber of the insert were removed, and a PBS-wet cotton-tipped applicator was used to wipe off all cells from the top membrane. Inserts were then fixed in 800 µL 4% PFA in PBS for 10 minutes at RT. Next, inserts were rinsed 3x with PBS and stained with 0.2% crystal violet (20% methanol/80% deionized water) for 5 to 10 minutes at RT in the dark. Inserts were washed three times with deionized water, dried, and then imaged. Cell counts were quantified and compared.

### RNA isolation and bulk mRNA sequencing

MDA-MB-231 breast cancer cells were seeded into 6-well plates with their respective media and allowed to adhere overnight. Once cells were at a confluence of 70-80%, they were washed with 1X DPBS and incubated with either pericyte-conditioned media or control media, with and without 10 nM of DTX or a vehicle control, for 24 hours. After treatment, total RNA was isolated using the RNeasy Mini Kit (Qiagen #74106) according to the manufacturer’s protocol. RNA was eluted in molecular biology-grade water. All RNA preparation and handling took place under RNase-free conditions. RNA quality and quantity were assessed using a NanoDrop spectrophotometer (Thermo Fisher Scientific, RRID:SCR_015804). RNA samples were stored at -20°C until further use.

Novogene performed library preparation and sequencing. Libraries were sequenced with an Illumina NovaSeq Z Plus in paired-end 150 bp mode, yielding 20-35 million read pairs per sample after quality control filtering. Paired-end reads were quantified against a decoy-aware GENCODE v45 (GRCh38) transcriptome index using Salmon v1.10.3 [12] with GC-bias, sequence-bias, and selective-alignment corrections. Transcript estimates were summarized to the gene level with *tximport* (length-scaled TPM; EnsDb.Hsapiens.v86 annotation) [13] and differential expression was tested in DESeq2 [14] retaining genes with a count ≥10 in at least four samples and applying ashr log-fold-change shrinkage. Genes with log_2_ fold-change > 1 and Benjamini-Hochberg FDR < 0.05 were called as differentially expressed [15]. Gene set enrichment was performed with clusterProfiler [16] against the MSigDB Hallmark collection [17] under default parameters, with genes ranked by sign (log_2_ fold-change) × −log_10_*P* and significance at FDR < 0.05. Pathway activity per sample was inferred with PROGENy [18] (top 500 footprint genes) using the multivariate linear model in decoupleR [19]. A SASP score per sample was computed as the mean row-scaled, variance-stabilized expression of 28 senescence-associated secretory genes, and groups were compared using a Wilcoxon rank-sum test.

We prioritized upstream ligands with NicheNet [20] (v2 human model). Receiver-expressed genes (baseMean ≥ 10) served as the background, and we considered only ligands with ≥50 predicted targets. The target sets were the CM-vs-control differentially expressed genes. Because DTX-CM vs. CM had few such genes, we used the 200 genes with the lowest P values for that contrast. For PCA, we removed experiment effects from a copy of the variance-stabilized counts with limma::removeBatchEffect [21], separately within each run. We used these corrected values for display only.

### Preparation of samples for metabolomics

Pericytes were seeded into 6-well plates in their respective media (ScienCell Research Laboratories #1201) with 2% FBS and 1% P/S. Once they reached approximately 80% confluency, the cells were washed with PBS and then incubated with 1 mL of the following: pericyte medium with 2% dialyzed FBS (Cytiva HyClone™ Fetal Bovine Serum (U.S.), Dialyzed #SH3007902), denoted as “untreated”; DMEM SFM; 0.2% DMSO (Vehicle); and 10 nM DTX treatment for 24 hours. Additionally, one well in the 6-well plate was reserved for cell counting to examine and compare the drugs’ effects on pericyte cell count after 24 hours of treatment. Samples were transferred to the Metabolomics Platform on dry ice and immediately stored at -80 °C.

### Polar metabolite profiling using liquid chromatography-mass spectrometry (LC-MS)

30 µL of cell culture medium was mixed with 120 µL of 75/25 acetonitrile/methanol containing 0.1% formic acid, vortexed, incubated at 4°C for 20 minutes, centrifuged at 18,000g for 20 minutes at 4°C, and the supernatant was transferred to an LC-MS vial. The metabolite separation was performed on a Thermo Scientific Vanquish Horizon UHPLC system using an iHILIC-(P) Classic (2.1×150 mm, 5 µm; part # 160.152.0520; HILICON AB) column. The high-resolution Orbitrap IQ-X/ID-X Tribrid mass spectrometer (Thermo Scientific) with a H-ESI probe operating in switch polarity was used to detect and quantify the metabolite levels. The mobile phase A(MPA) was 20 mM ammonium bicarbonate at pH 9.6, adjusted by ammonium hydroxide addition, and MPB was acetonitrile. The column temperature, injection volume, and flow rates were 40°C, 2 µL, and 0.2 mL/minute, respectively. The chromatographic gradient was 0 minutes: 85% B, 0.5 minutes: 85% B, 18 minutes: 20% B, 20 minutes: 20% B, 20.5 minutes: 85% B and 28 minutes: 85% B. MS parameters were as follows: Acquisition range of 70-1000 m/z at 60K resolution, spray voltage:3600V for positive ionization and 2800 for negative ionization modes, sheath gas: 35, auxiliary gas: 5, sweep gas: 1, ion transfer tube temperature: 250°C, vaporizer temperature: 350°C, AGC target: 100%, and a maximum injection time of 118 ms. Pooled QC samples were injected intermittently to assess detection reproducibility and metabolite stability over time. Data acquisition was done using the Xcalibur software (Thermo Scientific), and data analysis was performed using Compound Discoverer 3.3 (± 5 ppm) & Skyline software [22]. Metabolite identification was performed by matching retention time and MS/MS fragmentation to an in-house database generated from commercially available reference standards.

### Compass

We estimated reaction activity from the bulk RNA-seq data of MDA-MB-231 cells exposed to conditioned media from DTX-treated (DTX-CM) or vehicle-treated (CM) pericytes (n = 4 per group, N = 8) using Compass [23] with the Recon2.2 genome-scale metabolic model, run with default settings on the University of Chicago high-performance computing cluster. Compass is a flux balance analysis-based algorithm that uses gene expression to assign each reaction a penalty, which reflects how inconsistent the reaction’s activity is with the expression data. High penalties therefore mark reactions that are unlikely to be active. Compass reports relative reaction activity and does not estimate absolute flux rates. The negative log of the penalty is used as the reaction consistency score, so higher scores indicate greater predicted activity. Reactions with confidence values of 0 or 4 were retained, and reactions with penalty scores close to 0 were excluded. Reversible reactions are modeled as separate forward (_pos) and reverse (_neg) directions, and reactions with identical scores in all samples were removed, leaving 6,522 directional reactions. Reactions were assigned to subsystems with the Recon2.2 annotation distributed with Compass. We compared groups using Welch’s t-test on reaction penalties and calculated Cohen’s d as (mean DTX-CM minus mean CM) / pooled SD on consistency scores, so that positive values indicate higher predicted activity in DTX-CM. Benjamini-Hochberg FDR correction was applied to the p-values.

### Statistical analysis

Data were analyzed in GraphPad Prism 10 (Boston, MA) and PlotTwist 1.0 (DRIFTLabs). Results are presented as Mean ± SEM, as noted. P values were determined from biological, not technical, replicates, and against an alpha value of 0.05. Statistical tests are explicitly included in the results and Fig. legends.

## RESULTS

### Pericyte-secreted factors drive a proliferative, high-grade-like program in triple-negative breast cancer cells

To determine how the pericyte secretome shapes tumor-cell behavior, we cultured MDA-MB-231 triple-negative breast cancer cells in pericyte-conditioned media (CM) or control media and profiled them by RNA-sequencing. Differential expression analysis identified 40 genes, 23 up and 17 down (Fig. 1a, Supp. Fig. S1a, Supp. Table S1). Principal component analysis also separated the two conditions into non-overlapping clusters, indicating a reproducible and coordinated transcriptional response (Supp. Fig. S1b). The most strongly induced genes were nicotinamide N-methyltransferase (*NNMT*) and the matrix-remodeling inhibitor *SERPINE1/PAI-1* followed by the hypoxia- and angiogenesis-associated genes *ANGPTL4* and *DDIT4*. *NNMT* overexpression is a hallmark of aggressiveness across many cancers [24], and *SERPINE1/PAI-1* is a poor prognostic marker in breast cancer [25]. These markers indicate that the secretome shifts the cells toward a more aggressive transcriptional state.

**Fig. 1.**
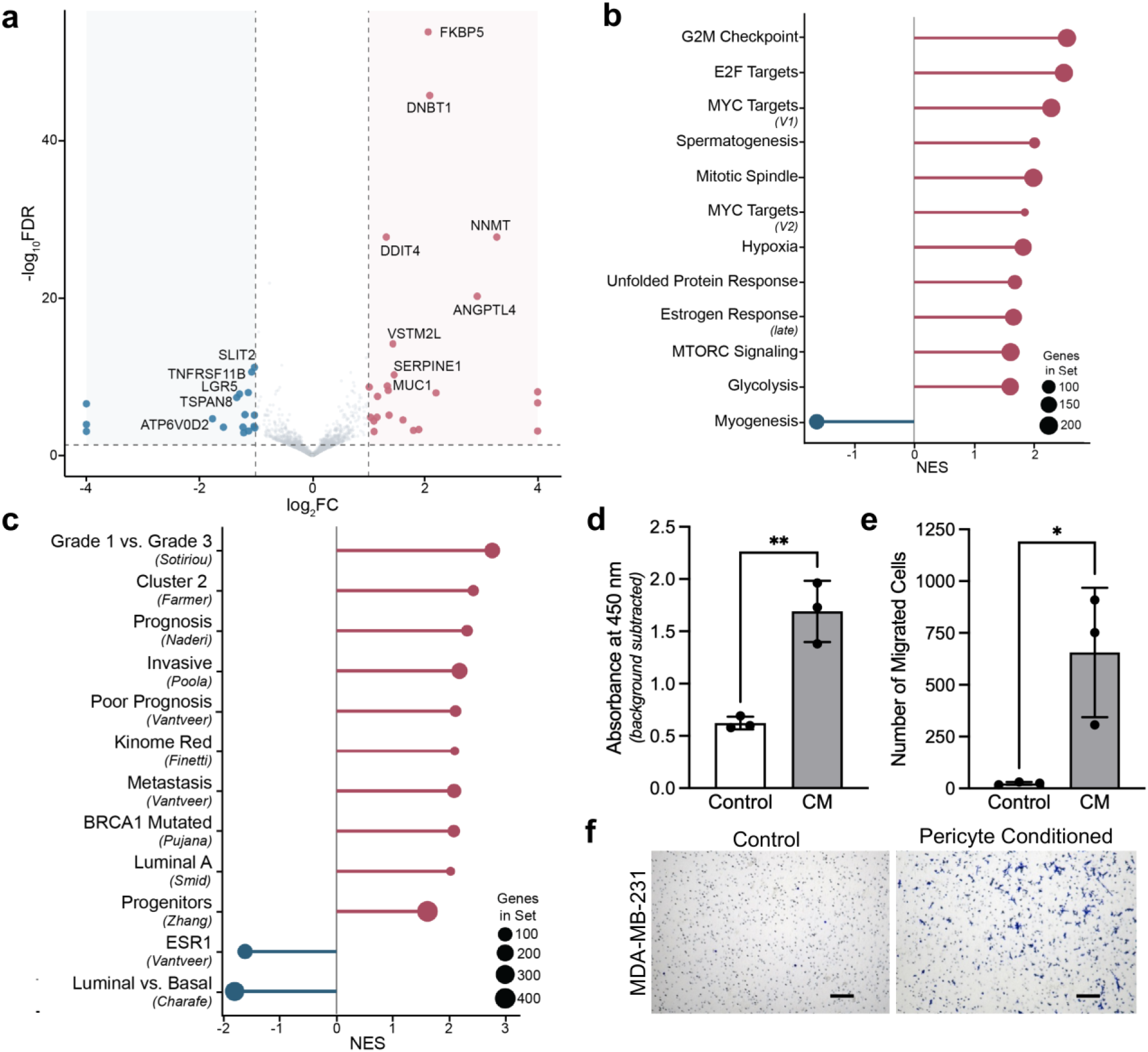
Pericyte conditioned media reprograms the TNBC transcriptome, driving proliferation. **(a)** Volcano plot of differential expression in MDA-MB-231 cells cultured in pericyte conditioned media (CM) versus control media, with selected genes labeled. Dashed lines mark |log₂FC| = 1 and FDR = 0.05. **(b)** Hallmark gene set enrichment for CM versus control media; the 12 most significant sets are shown. Position is the normalized enrichment score (red, enriched; blue, depleted) and dot size is the gene set size. **(c)** fgsea overlap with published human breast-cancer signatures (MSigDB C2:CGP, 23 significant at FDR < 0.05, top 12 shown). **(d)** Proliferation of MDA-MB-231 cells in control media or pericyte conditioned media, as percent proliferating cells relative to control. Mean ± SEM, n=3. **, p < 0.01. **(e)** Transwell migration of MDA-MB-231 cells toward control media or pericyte conditioned media, as fold change in migrated cells relative to control. Mean ± SEM, n=3. *, p < 0.05. **(f)** Representative images of MDA-MB-231 cells that migrated through the transwell membrane, control versus pericyte conditioned media. Scale bar, 200 µm.

To define the programs underlying this state, we performed Hallmark gene-set enrichment analysis. G2M Checkpoint (NES = 2.6), E2F Targets (NES = 2.5), MYC Targets V1 (NES = 2.3) and Mitotic Spindle (NES = 2.0) were the most enriched sets and together formed a dominant proliferation signature. Hypoxia (NES = 1.8), Glycolysis (NES = 1.6) and mTORC1 Signaling (NES = 1.6) were also enriched. In all, 20 of 50 Hallmark sets reached significance (FDR < 0.05, Fig. 1b). Given that this response must be driven by factors in the pericyte secretome, we used NicheNet ligand-activity inference to nominate candidate drivers. The top-ranked ligands were TGFB1, IL1B, LPA and FGF4. The top set also included the canonical perivascular ligand CXCL12/SDF-1 and TGFB2 (Supp. Fig. S1c). TGF-β is a primary driver of epithelial-mesenchymal transition and metastasis in breast cancer [26].

We next asked whether the CM-induced signature resembled aggressive human disease and found that it was enriched for programs of high histologic grade (Sotiriou Genomic Grade Index [27], proliferation [28], and poor prognosis [29]). The signature also carried a basal, non-luminal identity consistent with the triple-negative phenotype of the cells (Fig. 1c). This overlap is largely driven by the shared proliferative program. These data indicate that the pericyte secretome drives a proliferative program in triple-negative breast cancer cells that resemble high-grade, poor-prognosis human breast cancer.

To test whether this program translated into cancer cell behavior, we measured proliferation and migration in three breast cancer cell lines after 72 hours in CM or control media. MCF-7 and T-47D cells represent ER+ breast cancer and MDA-MB-231 cells represent triple-negative breast cancer. CM increased proliferation of all three lines (p < 0.01 in MDA-MB-231, Fig. 1d, and p < 0.05 in MCF-7 and T-47D, Supp. Fig. S2a, S2b). CM also increased Transwell migration (p < 0.05, Fig. 1e, 1f, Supp. Fig. S2c-f). Together, these data indicate that the pericyte secretome promotes both growth and motility. The migratory response suggests that a secreted chemoattractant is involved, consistent with the role of CXCL12/SDF-1, one of our top-ranked ligands, in breast cancer chemotaxis and metastasis [30]. The cells also expressed receptors for several top-ranked ligands, including IL1R1 for IL1B and TGFBR2 for TGFB1 and TGFB2, both increased in CM (FDR < 0.05, Supp. Fig. S1d). Together, these data suggest that the pericyte secretome gives breast cancer cells the ligands to grow and migrate, and that the cells carry the receptors to respond.

### Docetaxel enriches pyrimidine nucleosides in the pericyte secretome without remodeling it globally

To identify components that could account for these effects, we performed untargeted polar metabolomics on pericyte-conditioned media. Vehicle-treated and DTX-treated pericytes showed broadly conserved patterns of metabolite production and consumption. In vehicle-treated pericytes, guanine, cytosine, lactate, hypoxanthine, uridine, and proline were among the most prominently secreted metabolites (Fig. 2a). Conversely, guanosine, pyruvate, inosine, and adenine were depleted relative to blank media, consistent with net consumption. Docetaxel treatment retained this overall pattern, with cytidine and thymidine becoming more prominent, and the same four metabolites remaining depleted (Fig. 2b). To resolve what docetaxel changed, we compared the two conditions directly. Several metabolites were significantly increased after docetaxel treatment, including guanine, uridine, proline, thymidine, cytidine, oxoproline (pyroglutamate), inosine and N,N,N-trimethyllysine (p < 0.05, Fig. 2c). Guanine and uridine showed the strongest combined effect size and significance.

**Fig. 2.**
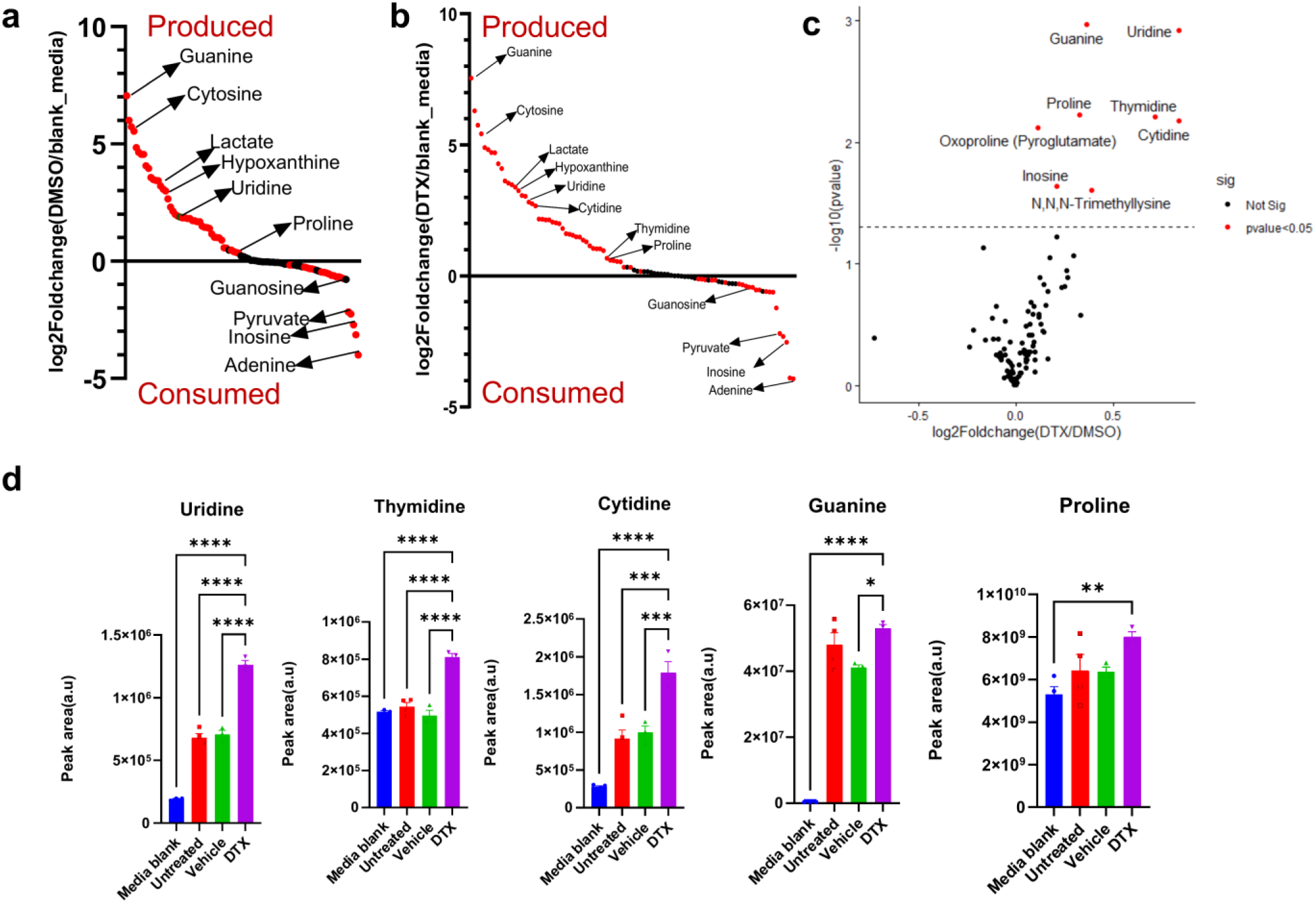
Docetaxel alters the extracellular metabolite profile of pericytes compared with DMSO. (**a,b**) Ranked log2 fold-change values for metabolites detected in conditioned media from (**a**) DMSO/vehicle-treated pericytes or (**b**) docetaxel-treated pericytes relative to blank media. Positive values indicate metabolites produced/secreted into the media, whereas negative values indicate metabolites consumed from the media. **(c)** Volcano plot showing differential extracellular metabolites in docetaxel-treated versus DMSO/vehicle-treated conditioned media. Red points denote significantly altered metabolites (p < 0.05). **(d)** Peak areas (a.u.) for uridine, thymidine, cytidine, guanine, and proline measured in media blank, untreated, vehicle, and docetaxel (DTX) conditioned media derived from pericytes. Mean ± SEM shown. 3-4 technical replicates analyzed. Ordinary one-way ANOVA; Dunnett’s multiple comparison test compared to DTX. a.u. = arbitrary units *p < 0.05, **p < 0.01, ***p < 0.001, ****p < 0.0001

We next confirmed these findings by targeted profiling with media blank, untreated, and vehicle-treated controls (Fig. 2d). Uridine and thymidine peak areas were higher in docetaxel-treated samples than in all three control groups (p < 0.0001). Cytidine was also highest in the docetaxel group (p < 0.0001 vs. blank media, p < 0.001 vs. untreated and vehicle). Guanine was increased relative to the media blank (p < 0.0001) and modestly relative to vehicle (p < 0.05), but not relative to untreated pericytes. Proline was increased only relative to the media blank (p < 0.01), a weaker effect than the untargeted comparison suggested.

Docetaxel changed the metabolic environment quantitatively rather than globally. The most robust effects were on the pyrimidine nucleosides uridine, thymidine, and cytidine. These measurements reflect extracellular abundance and cannot distinguish increased release from reduced uptake, cell death or altered turnover. Given the effects of conditioned media on proliferation and migration, these nucleosides are candidate contributors to a supportive microenvironment.

### Docetaxel-conditioned pericyte media adds a SASP-like inflammatory program to the cancer cell response and makes its net effect growth-inhibitory

Pericytes coexist with cancer cells within the microenvironment and are therefore exposed to chemotherapy. We asked how docetaxel alters the pericyte secretome by profiling MDA-MB-231 triple-negative breast cancer cells exposed to conditioned media from DTX-treated or vehicle-treated pericytes. At the global level the two responses were nearly indistinguishable, and principal component analysis did not separate them (Supp. Fig. S3a, Supp. Table S2). Only LOC389831, an uncharacterized locus, passed the stringent threshold (log₂FC > 1; Supp. Fig. S3b). The DTX-CM response also remained highly concordant with the CM program, with a log₂FC correlation of r = 0.91 versus control media (Supp. Fig. S3c). Global expression therefore did not capture the change, so we turned to the pathway level to resolve it.

Hallmark gene-set enrichment identified 16 of 50 sets as significant. This result and an orthogonal cytokine-signaling footprint were dominated by TNFα-NFκB signaling (Supp. Fig. S3d, S3e). Inflammatory response, IL6-JAK-STAT3, KRAS, protein secretion, and epithelial-mesenchymal transition were also enriched. To place these changes relative to the basal CM program, we mapped hallmark enrichment in CM versus media against DTX-CM (drug-CM) versus CM and three programs emerged (Fig. 3a, 3b). The first group comprised pathways amplified by docetaxel conditioning, including TNFα-NFκB, Hypoxia, and Glycolysis. The second comprised drug-specific pathways not driven by CM alone, including KRAS, Inflammatory Response, IL6-JAK-STAT3, Protein Secretion, and EMT. The third comprised CM-only pathways, namely the entire proliferation module of G2M, E2F and MYC, which the drug left unchanged. PROGENy pathway-activity inference independently confirmed rising TNFα and NFκB activity from CM to DTX-CM (Supp. Fig. S3f).

**Fig. 3.**
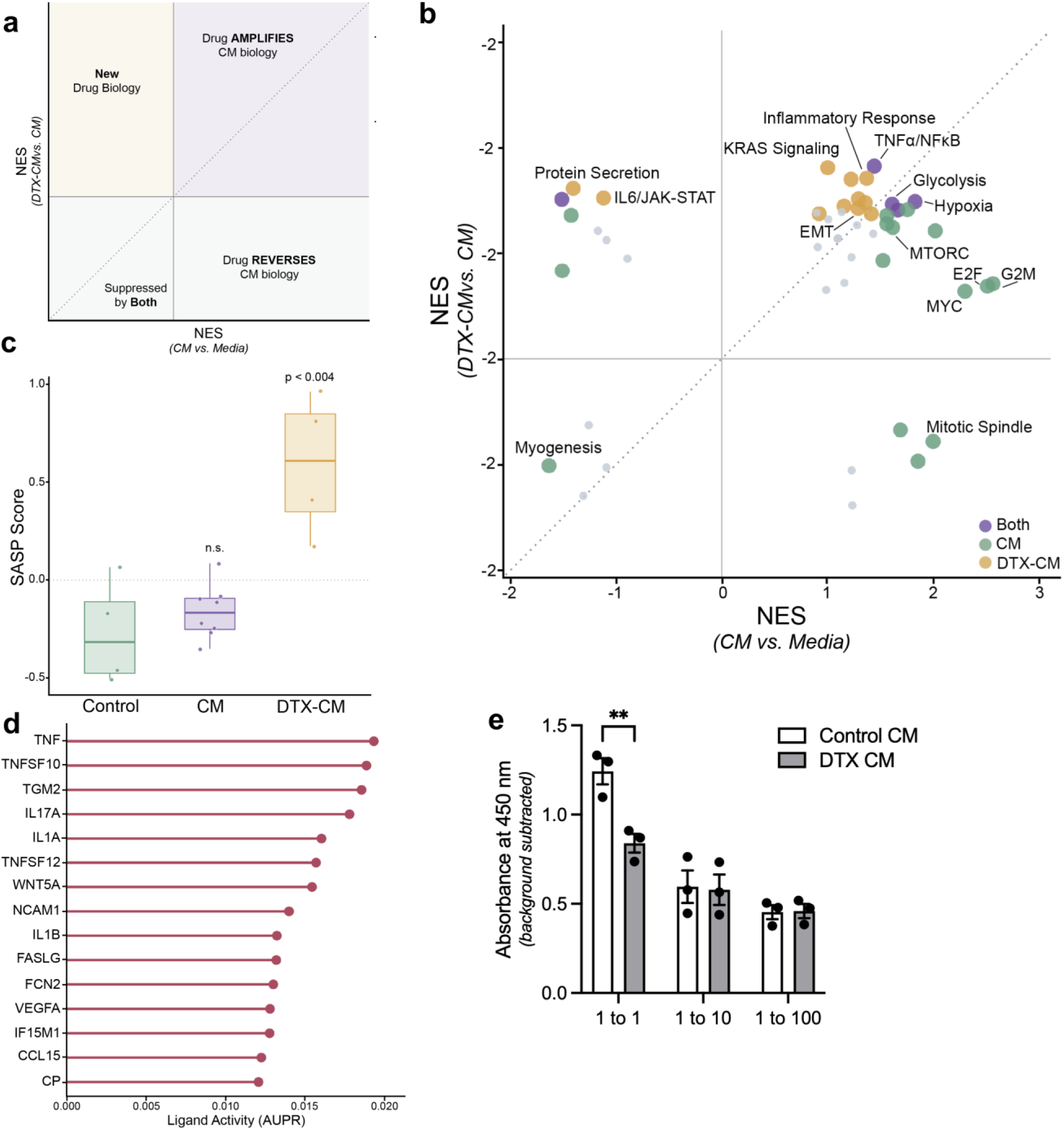
Docetaxel-conditioned pericyte media adds an inflammatory, SASP-like program to the secretome response. **(a)** Schematic for reading **(b)**. Upper right, CM pathway amplified by docetaxel; upper left, new drug-only biology; lower right, CM pathway reversed by docetaxel; lower left, suppressed in both. No data plotted. **(b)** Hallmark pathway map. NES for CM vs media (x) against NES for DTX-CM vs CM (y), 50 sets. Points colored by significance: both (purple, n = 5), drug-only (yellow, n = 11), CM-only (green, n = 15), neither (grey, n = 19). **(c)** SASP score per library, the mean row z-score across 28 canonical SASP genes; points are individual libraries. The P value is a two-sided Wilcoxon test of DTX-CM versus CM, **, p < 0.01. **(d)** NicheNet ligand-activity ranking, DTX-CM versus CM, top 15 of 955 ligands by corrected AUPR. **(e)** Proliferation of MDA-MB-231 cells in vehicle CM or docetaxel-conditioned CM (DTX-CM), across a 1:1, 1:10 and 1:100 dilution series. Mean ± SEM, n=3. **, p < 0.01.

Given that this drug-specific program centered on inflammatory signaling, we asked whether it resembled a senescence-associated secretory phenotype (SASP), the inflammatory secretory program through which senescent cells drive paracrine invasion and malignancy [31]. The SASP score was elevated specifically in DTX-CM (p = 0.004), but not in vehicle CM (Fig. 3c). The leading-edge genes (the core genes driving the set enrichment) were classic NFκB and chemokine outputs, led by the chemokines CXCL8 and CXCL1/3 (Supp. Fig. S3g). NicheNet traced this to TNF-superfamily and IL-1/IL-17-family ligands, including TNF, IL1B, and TNFSF10/TRAIL (Fig. 3d). This response is consistent with our earlier finding of IL-6 enrichment in the docetaxel-pericyte secretome [11].

Together, these data suggest that the docetaxel conditioning did not overwrite the secretome response but added to it. The DTX-CM secretome is predicted to carry a TNF/IL-1/IL-6 program that engages NFκB and STAT3 signaling in the cancer cell, an axis that supports an aggressive phenotype in triple-negative breast cancer [32]. This pattern mirrors therapy-induced senescence, in which chemotherapy drives a SASP in non-malignant cells that promotes inflammation and relapse [33].

To test the functional consequences of this secretome, proliferation was measured in three breast cancer cell lines (MDA-MB-231, MCF-7, and T-47D) across a dilution series of DTX-CM or vehicle-CM. At the undiluted dose, docetaxel-CM reduced proliferation of all three lines relative to vehicle-CM (Fig. 3e, Supp. Fig. S4a, S4b). The effect was dose-dependent and lost at 1:10 and 1:100 dilutions, consistent with a secreted factor. To rule out carryover of docetaxel into the conditioned media and to ask whether pericyte CM protects cancer cells from the drug, we generated docetaxel dose-response curves in control CM or pericyte CM. The docetaxel IC_50_ did not differ between the media conditions (Supp. Fig. S4c-e). Thus, pericyte CM did not measurably alter docetaxel sensitivity in this assay, and the growth inhibition by DTX-CM was not explained by residual drug.

Taken together, docetaxel conditioning shifted the net effect of the pericyte secretome from growth-promoting to growth-inhibitory at the highest concentration. The SASP-like program fits this shift, since it includes the pro-apoptotic ligands TNFSF10/TRAIL and FASLG, which NicheNet ranked among the top candidate ligands.

### Metabolic modeling predicts higher transport and sterol metabolism and lower nucleotide interconversion in cancer cells exposed to DTX-CM

To ask how the secretome rewires recipient-cell metabolism, we applied Compass, which scores the potential activity of each metabolic reaction from gene expression [23], to the RNA-seq profiles of MDA-MB-231 cells exposed to CM or DTX-CM. The reactions that changed most fell into three groups: transport, sterol and nucleotide metabolism (Fig. 4a). Of 6,522 modeled reactions, 59 (0.9%) differed at a nominal p < 0.05, with 37 predicted higher and 22 predicted lower in DTX-CM (Fig. 4b). Because no reaction passed FDR correction, we ranked reactions by effect size and followed the largest effects (Fig. 4c).

**Fig. 4.**
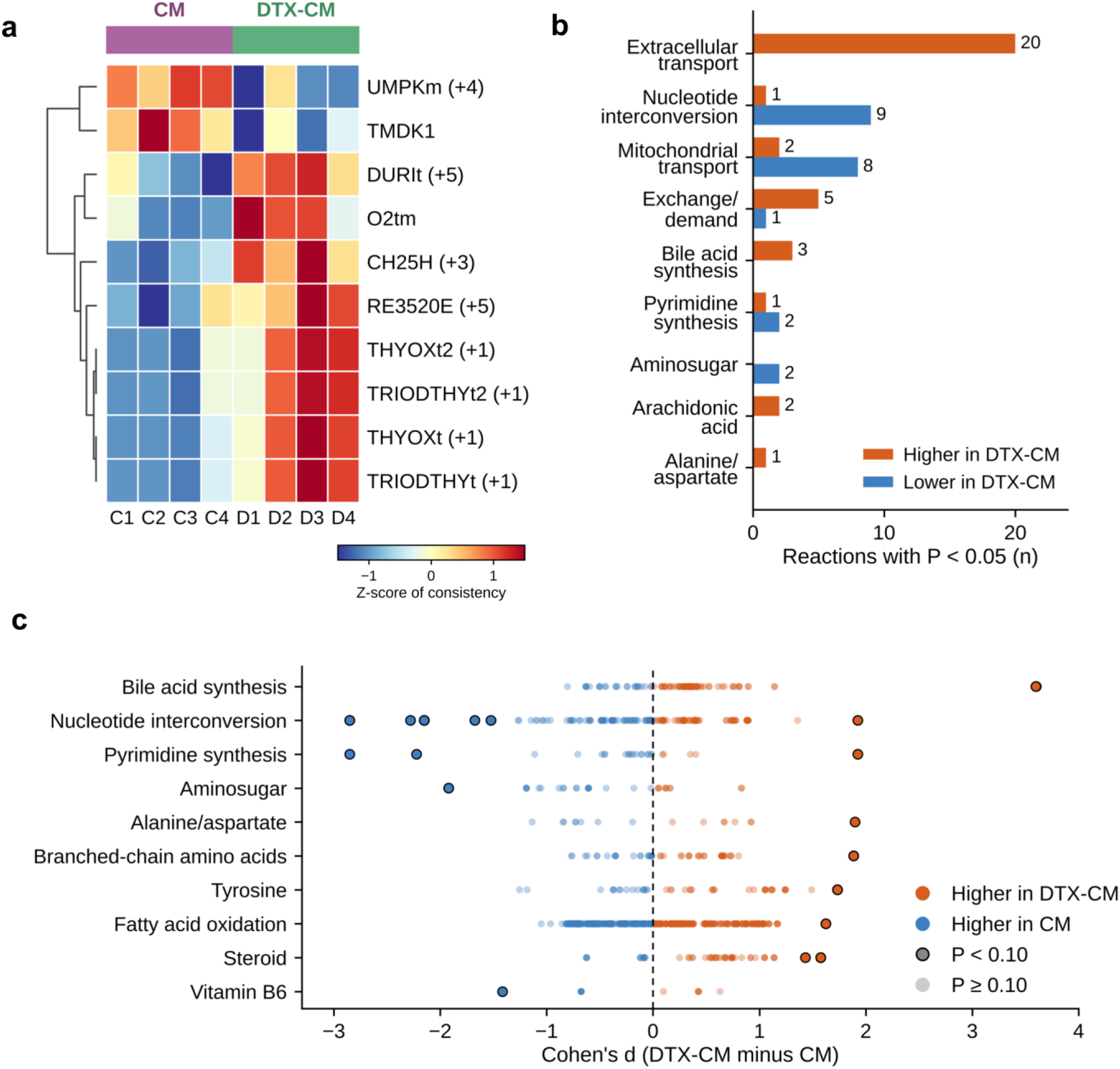
Metabolic modeling predicts higher extracellular transport and lower nucleotide interconversion in cancer cells exposed to DTX-PCM. **(a)** Reaction consistency heatmap. Z-score of the Compass consistency score per reaction (higher = more active) for the 10 most significant distinct reaction profiles, clustered by reaction. Coupled reactions with identical profiles are collapsed into one row, with the number of additional reactions in parentheses (31 reactions in total). Columns are individual samples (C1-C4, Veh-PCM. D1-D4, DTX-PCM). **(b)** Reactions per subsystem with p < 0.05 (Welch’s t-test), DTX-PCM versus Veh-PCM, top 9 subsystems. Orange, higher in DTX-PCM (n = 37 reactions overall). Blue, lower in DTX-PCM (n = 22). No reaction passed FDR correction (q < 0.05) among 6,522 modeled reactions. **(c)** Effect size per reaction. Cohen’s d (DTX-PCM minus Veh-PCM) by subsystem, 10 subsystems. Points are individual reactions, colored by the group with higher predicted activity (orange, DTX-PCM. Blue, Veh-PCM). Opaque, outlined points, p < 0.10. Faded points, p ≥ 0.10. MDA-MB-231 cells, n = 4 per group.

Transport was the clearest signal. All 20 significant extracellular transport reactions were higher in DTX-CM (Fig. 4b), including routes for thyroid hormone (THYOXt and TRIODTHYt) and nucleosides (DURIt, DCYTt and DADNt4). Sterol metabolism carried the largest effect sizes. CH25H, which converts cholesterol to 25-hydroxycholesterol, reached d = +3.60 (p = 0.004), as did RE2814R and r0672 (Fig. 4a, 4c). 25-hydroxycholesterol activates liver X receptors and controls cholesterol efflux and inflammatory gene expression [34], which offers one link between this sterol shift and the inflammatory program in DTX-CM.

Nucleotide metabolism moved the other way. Nine of 10 significant nucleotide interconversion reactions were lower in DTX-CM, including mitochondrial UMP kinase (UMPKm, d = -2.85, p = 0.015) and thymidine kinase (TMDK1, d = -2.28, p = 0.020) (Fig. 4a, 4c). Nucleoside diphosphate kinases keep nucleotide pools, genome stability and growth in balance f [35]. Mitochondrial transport leaned lower as well, with 8 of 10 significant reactions reduced (Fig. 4b). In parallel, arachidonic acid metabolism rose in DTX-CM (Fig. 4a, 4b), led by RE3520E (d = +2.20, p = 0.022). Arachidonic acid feeds prostaglandin E2, which promotes tumor progression and shapes tumor immunity [36]. Together, these data indicate that DTX-CM pushes recipient cells toward greater hormone and nucleoside transport and oxysterol and eicosanoid production, while turning down mitochondrial nucleotide interconversion.

## DISCUSSION

Our study shows that the pericyte secretome supports breast cancer cell proliferation and migration, and that docetaxel reprograms it toward an inflammatory, SASP-like output. Pericyte-conditioned media (CM) induced a proliferative, high-grade-like program in cancer cells, marked by G2M, E2F and MYC signatures and strong induction of *NNMT* and *SERPINE1/PAI-1* [24, 25, 27–29]. Inhibition of *NNMT* in pericytes has been demonstrated recently in murine colorectal cancer to restore antitumor function of CD8+ T cells for cancer immunotherapy [37]. After docetaxel exposure, the pericyte secretome induced an inflammatory, SASP-like response in cancer cells, became enriched for pyrimidine nucleosides, and inhibited cell growth at high concentrations. Importantly, these findings align with studies showing that pericytes act as signaling cells that support tumor growth through secreted factors, including metabolites [4, 5, 10].

The docetaxel-induced program fits a model in which chemotherapy drives a SASP in non-malignant cells that feeds back on the tumor [31, 33, 38]. Pathway analysis showed a shift toward TNFα-NFκB, inflammatory response and IL6-JAK-STAT3 programs, and the SASP score was elevated specifically in cells exposed to DTX-CM, consistent with our earlier observation of IL-6 enrichment in the DTX-treated pericyte secretome [11]. Taxane-induced stromal injury also promotes breast cancer dormancy escape [39]. We propose that docetaxel does not switch pericyte support on or off but adds an inflammatory component to a growth-supporting secretome [40, 41]. Consistent with this, pericyte CM did not alter docetaxel potency, while DTX-CM inhibited growth at high concentrations. The nucleosides uridine, thymidine and cytidine, which accumulated in DTX-treated pericyte media, provide one candidate route, since stromal nucleoside release alters chemotherapy response in pancreatic cancer [42, 43]. Exploratory metabolic modeling of recipient cells predicted higher extracellular transport and lower mitochondrial nucleotide interconversion, which points to nucleoside handling as a candidate axis. Because these cells were not exposed directly to docetaxel, these changes are most likely mediated by pericyte-derived factors.

More broadly, our results support a model in which chemotherapy acts on the tumor twice, directly on cancer cells and indirectly through the pericytes that surround them. Pericyte-derived inflammatory signals may be targetable, as our earlier finding that IL-6 receptor blockade enhances docetaxel efficacy in ER+ breast cancer indicates [11]. By resolving how docetaxel reprograms the pericyte secretome, our work reframes the perivascular niche as an active participant in the response to chemotherapy, and the perivascular niche may be a new therapeutic target for improving taxane-based treatment.

## Conclusions

The pericyte secretome promotes proliferation and migration, and docetaxel reprograms it toward an inflammatory, SASP-like output enriched in pyrimidine nucleosides. These findings suggest that chemotherapy-exposed pericytes can reshape the tumor niche, and that pericyte-derived nucleosides and inflammatory signals are candidate therapeutic targets. Future work will identify the factors responsible and test their role in more physiologic settings.

## Data and code availability

Research outputs will be preserved and shared with the public via Knowledge@UChicago, the University of Chicago institutional repository. Knowledge@UChicago is managed by the University Library and uses a cloud-based hosted repository service called TIND as its platform. TIND is based on the Invenio software originally developed at the research institute CERN to manage its own scholarly output. TIND uses an OAIS-compliant approach to preservation, with redundancy backup, conversion to archival file formats, storage of archival packages on geographically separated servers, and fixity checking. To support discoverability and citation, DOIs are minted for deposits made to Knowledge@UChicago through the DataCite Fabrica service.

All data generated or analyzed during this study are included in the Article or as Supplemental Information. The source data underlying selected Fig.s and Supplementary Fig.s are provided as a Source Data file. The data underlying the findings of this study can be obtained from the corresponding author upon reasonable request. Please contact Samantha S. Yee at.

Data used in this study can be shared by the corresponding author upon reasonable request.

## Contributions

Conceptualization: K.M.F., M.H.C., S.S.Y.; Methodology: K.M.F., M.H.C., and S.S.Y.; Investigation: K.M.F., H.S., S.S.Y., N.A.J.; Formal analysis: K.M.F., M.H.C., H.S., S.S.Y.; Writing—original draft: K.M.F., S.S.Y., M.H.C., H.S., N.A.J.; Writing—review and editing: K.M.F., S.S.Y., M.H.C. All authors reviewed and approved the manuscript – K.M.F., S.S.Y., M.H.C., H.S., N.A.J., S.L.H. Funding Acquisition: K.M.F. and S.S.Y.; Supervision: S.S.Y.; Project administration: S.S.Y.

## Acknowledgements

We would like to thank Jinyang Li and Anthony Kossiakoff for cell lines. We appreciate the Metabolomics Platform at the University of Chicago Medicine Comprehensive Cancer Center (Metabolomics Platform Director: Hardik Shah). Also, we thank Jessica J. Kandel (University of Chicago) for her continuous support. Lastly, we thank Simon Schwörer (University of Chicago) for scientific discussion and help with submitting bulk mRNA-sequencing samples to Novogene.

## Ethics declarations

The authors declare no conflict of interests.

---

**Supplementary Figure S1.**
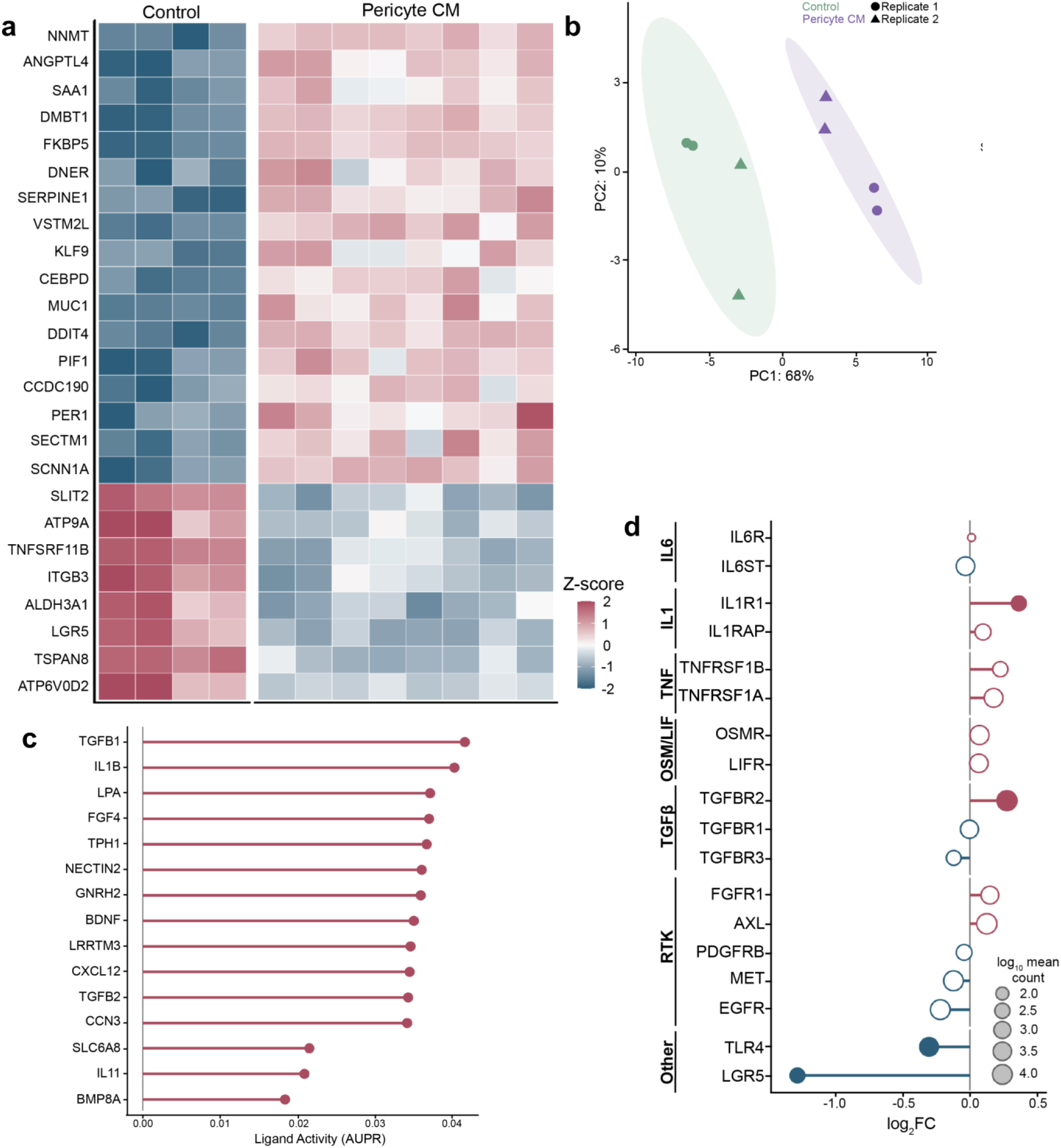
**(a)** Row z-scored expression heatmap of the top 25 differentially expressed genes (ranked by FDR among genes with mean normalized count ≥ 50) across control and conditioned-media libraries. **(b)** Principal component analysis of the media sequencing run (n = 8; 4 control, 4 CM) after removing the replicate covariate. Control media, green; pericyte conditioned media, purple; the two biological replicates are circles and triangles. Ellipses are 85% confidence regions. **(c)** NicheNet candidate ligands driving the CM response (top 15 of 955 candidates, by corrected AUPR). **(d)** Lollipop of 18 cytokine, growth-factor and related receptors grouped by family. Position is the log_2_ fold change in CM versus control media, dot size is the log_10_ mean normalized count (receptor abundance), filled dots are significant at FDR < 0.05 and open dots are not.

**Supplementary Figure S2.**
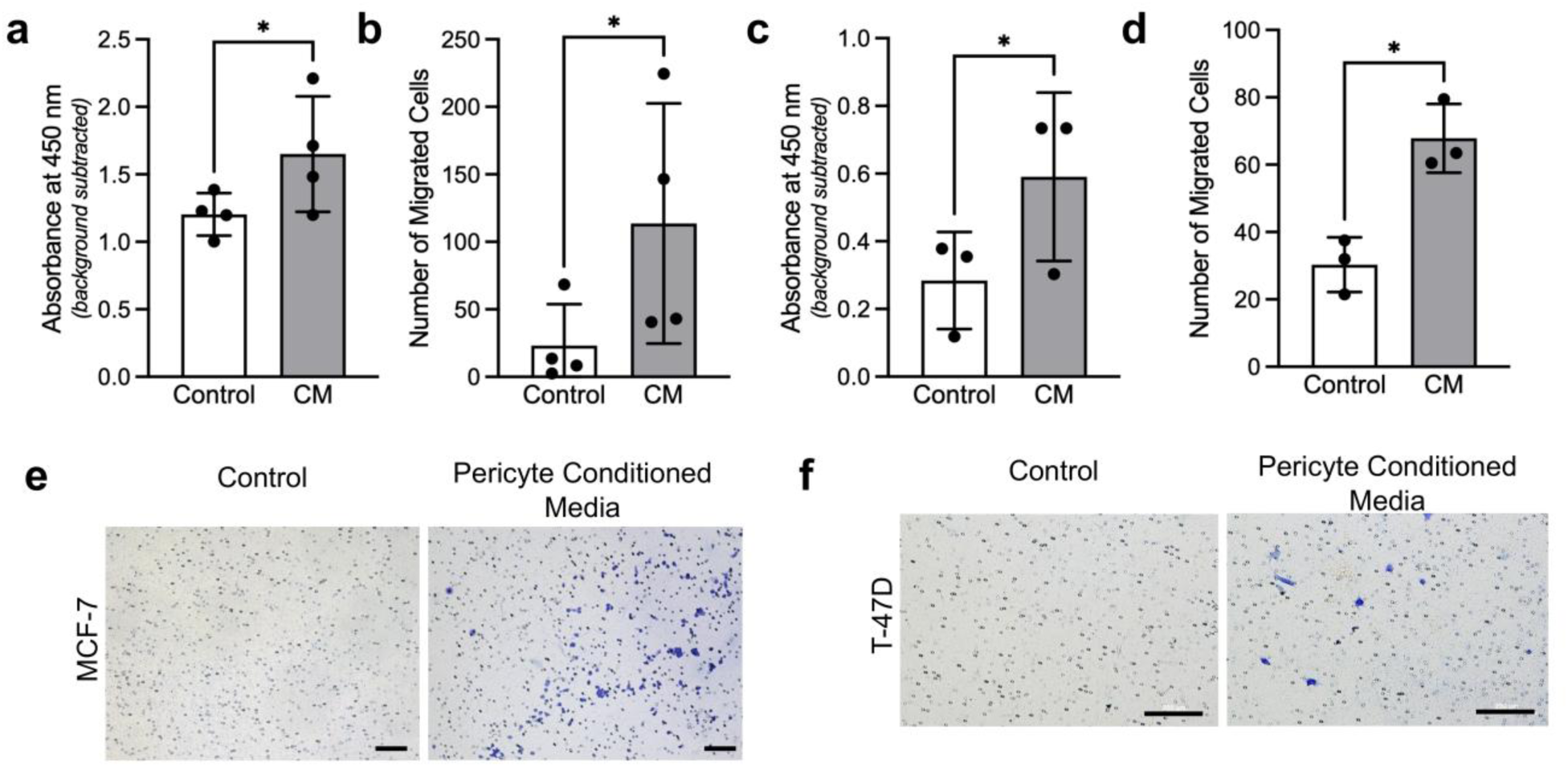
**(a, b)** Proliferation of MCF-7 **(a)** and T-47D **(b)** cells in control media or pericyte conditioned media, as percent proliferating cells relative to control. Mean ± SEM, n = 3. *, p < 0.05. **(c, d)** Transwell migration of MCF-7 **(c)** and T-47D **(d)** cells toward control media or pericyte conditioned media, as fold change in migrated cells relative to control. Mean ± SEM, n = 3-4. *, p < 0.05. **(e, f)** Representative transwell images of migrated MCF-7 **(e)** and T-47D **(f)** cells, control versus pericyte conditioned media. Scale bar, 200 µm.

**Supplementary Figure S3.**
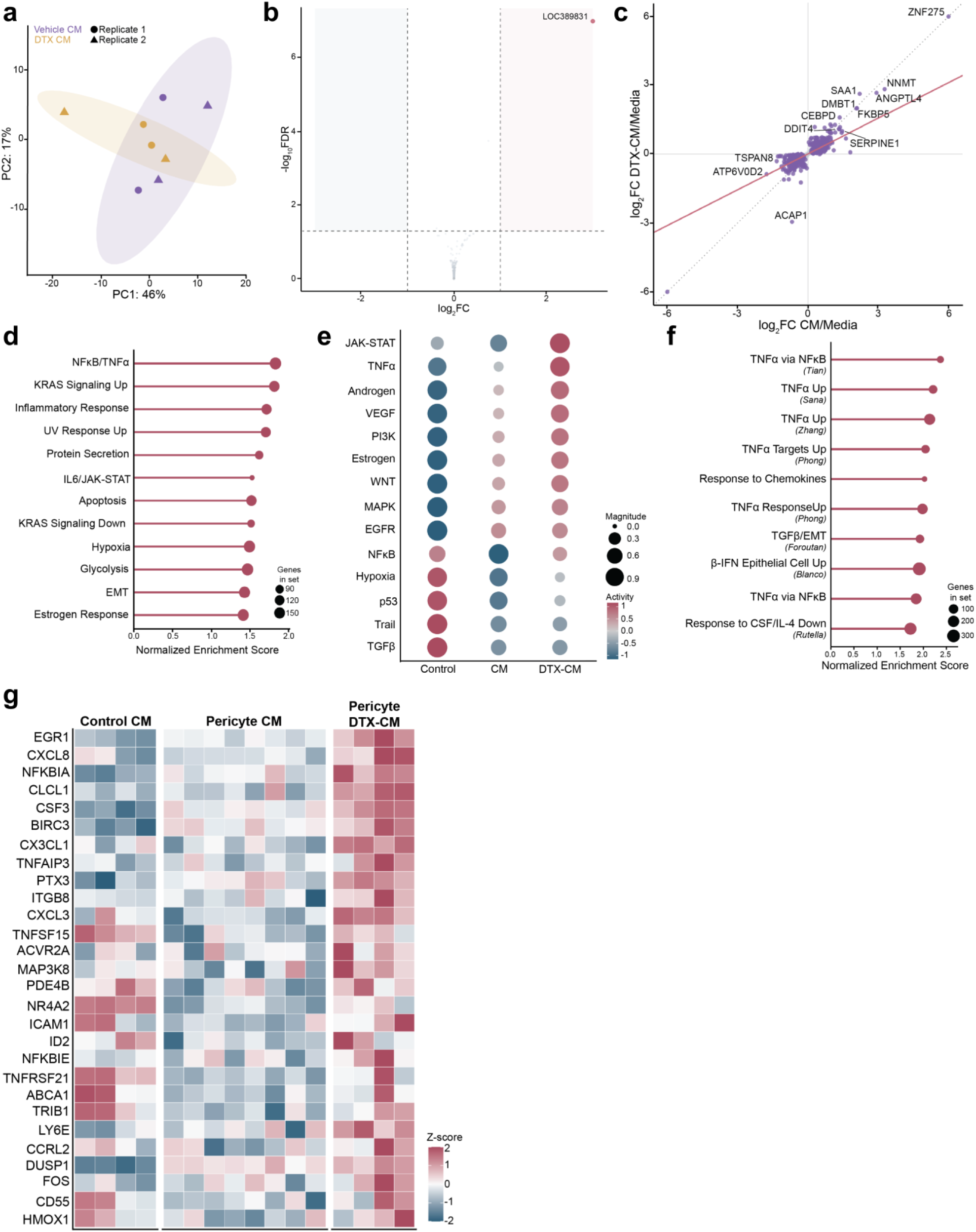
**(a)** PCA of control CM (purple) and docetaxel-CM (yellow) does not separate conditions. **(b)** Volcano plot of DTX-CM versus vehicle CM. Only LOC389831 passes | log_2_FC | > 1 and FDR < 0.05. **(c)** log_2_FC concordance, CM (x) versus DTX-CM (y) relative to control media, for 12,784 genes with mean count ≥ 30. Pearson r = 0.91. **(d)** Hallmark GSEA of DTX-CM versus CM. Top 12 sets shown, 16 of 50 significant at FDR < 0.05. **(e)** Cytokine-signaling footprint against 304 curated pathways. Top 10 shown, 25 significant at FDR < 0.05 versus 6 for CM alone. **(f)** PROGENy activity across 14 pathways, condition means row z-scored. **(g)** Row z-scored expression of 28 leading-edge genes from the TNFα-NFκB, IL6-JAK-STAT3 and inflammatory-response Hallmark sets, ranked by log_2_FC.

**Supplementary Figure S4.**
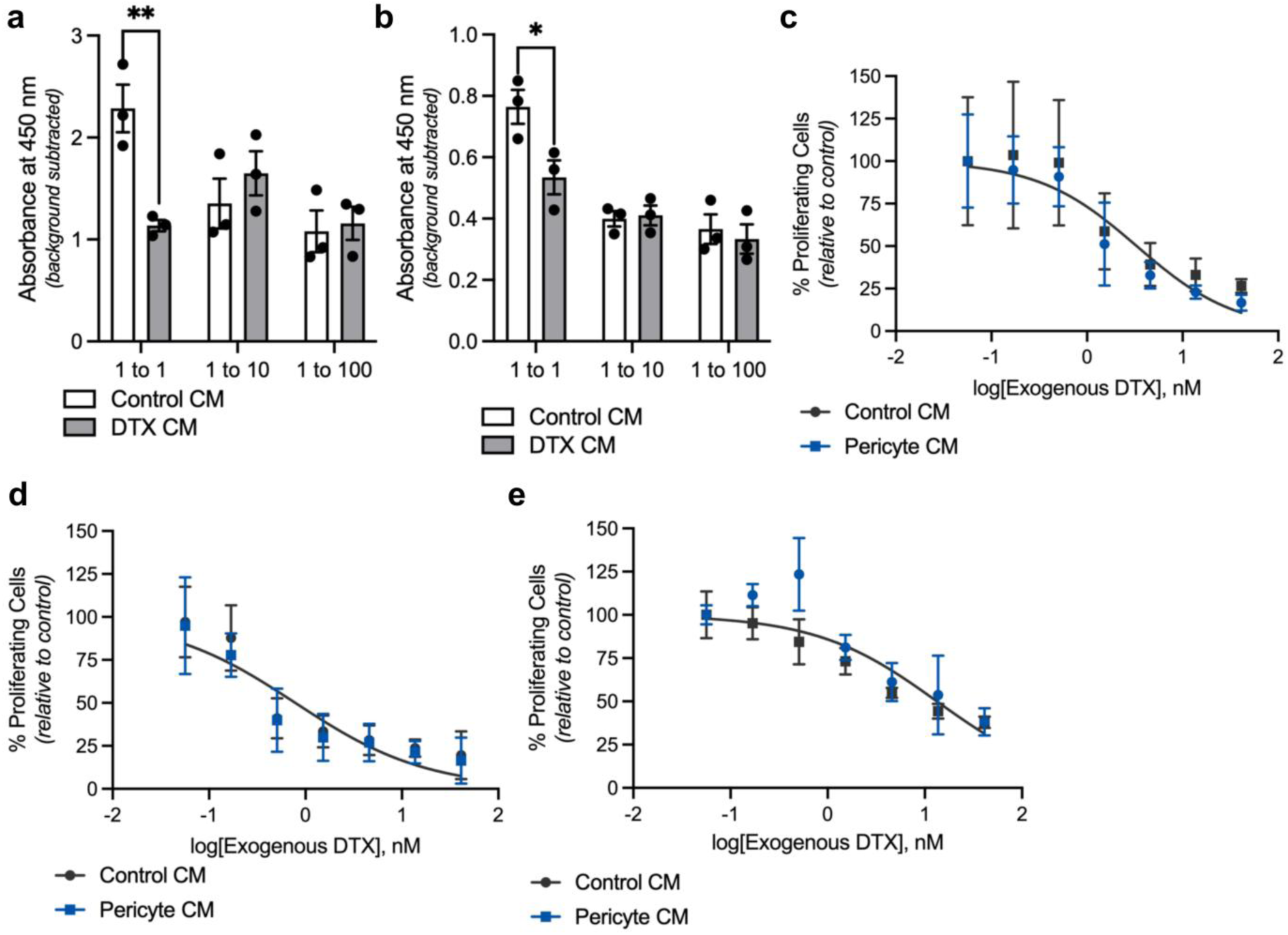
**(a, b)** Proliferation of MCF-7 **(a)** and T-47D **(b)** cells in control CM or DTX CM, across a 1:1, 1:10 and 1:100 dilution series. Mean ± SEM, n = 3. DTX CM reduced proliferation at the undiluted dose (MCF-7 **, p < 0.01; T-47D *, p < 0.05). **(c-e)** Docetaxel dose-response in MDA-MB-231 **(c)**, MCF-7 **(d)**, and T-47D **(e)** in control CM or pericyte CM normalized to DMSO. IC_50_ values did not differ significantly between media conditions in any cell line (MDA-MB-231, IC_50_ = 4.46 vs. 2.51 nM, p = 0.56; MCF-7, IC_50_ = 0.90 vs. 0.68 nM, p = 0.79; T-47D, IC_50_ = 10.29 vs. 16.18 nM, p = 0.064, extra sum-of-squares F test), n = 3-4.

## REFERENCES

1. Harris, M.A., et al., Towards targeting the breast cancer immune microenvironment. Nat Rev Cancer, 2024. 24(8): p. 554–577.

2. Postigo-Corrales, F., et al., Docetaxel Resistance in Breast Cancer: Current Insights and Future Directions. Int J Mol Sci, 2025. 26(15).

3. National Cancer, I., Surveillance, Epidemiology, and End Results Program. Cancer Stat Facts: Female Breast Cancer.

4. Zhang, C., et al., Methionine secreted by tumor-associated pericytes supports cancer stem cells in clear cell renal carcinoma. Cell Metab, 2024. 36(4): p. 778–792.e10.

5. Jordan, A.M., R.J.G. Drake, and K.M. Hodivala-Dilke, Angiocrine and pericrine signaling: how endothelial cells and pericytes drive cancer progression and therapy resistance. Physiol Rev, 2026. 106(1): p. 87–119.

6. Hosaka, K., et al., Pericyte-fibroblast transition promotes tumor growth and metastasis. Proc Natl Acad Sci U S A, 2016. 113(38): p. E5618–27.

7. Liu, Y., et al., Single-cell and spatial transcriptome analysis of breast cancer tumor-associated fibroblast heterogeneity and its mediated remodeling of the tumor microenvironment. BMC Cancer, 2026. 26(1).

8. Ismailov, A. and M. Poptsova, Pericyte-derived cancer-associated fibroblasts correlate with poor survival and are enriched after chemoradiotherapy in glioblastoma. PLoS One, 2026. 21(9): p. e0355902.

9. Safarzad, M., Pericyte contact alters endothelial cell metabolism by promoting exchange of lactate through SLC16A3, in bioRxiv. 2025.

10. Gaceb, A. and G. Paul, Pericyte Secretome. Adv Exp Med Biol, 2018. 1109: p. 139–163.

11. Przanowska, R.K., et al., IL-6R blockade with tocilizumab disrupts pericyte- and tumor cell-driven IL-6/STAT3 signaling, enhancing docetaxel efficacy in ER+ breast cancer. Scientific Reports, 2026.

12. Patro, R., et al., Salmon provides fast and bias-aware quantification of transcript expression. Nat Methods, 2017. 14(4): p. 417–419.

13. Soneson, C., M.I. Love, and M.D. Robinson, Differential analyses for RNA-seq: transcript-level estimates improve gene-level inferences. F1000Res, 2015. 4: p. 1521.

14. Love, M.I., W. Huber, and S. Anders, Moderated estimation of fold change and dispersion for RNA-seq data with DESeq2. Genome Biol, 2014. 15(12): p. 550.

15. Stephens, M., False discovery rates: a new deal. Biostatistics, 2017. 18(2): p. 275–294.

16. Wu, T., et al., clusterProfiler 4.0: A universal enrichment tool for interpreting omics data. Innovation (Camb), 2021. 2(3): p. 100141.

17. Liberzon, A., et al., The Molecular Signatures Database (MSigDB) hallmark gene set collection. Cell Syst, 2015. 1(6): p. 417–425.

18. Schubert, M., et al., Perturbation-response genes reveal signaling footprints in cancer gene expression. Nat Commun, 2018. 9(1): p. 20.

19. Badia, I.M.P., et al., decoupleR: ensemble of computational methods to infer biological activities from omics data. Bioinform Adv, 2022. 2(1): p. vbac016.

20. Browaeys, R., W. Saelens, and Y. Saeys, NicheNet: modeling intercellular communication by linking ligands to target genes. Nat Methods, 2020. 17(2): p. 159–162.

21. Ritchie, M.E., et al., limma powers differential expression analyses for RNA-sequencing and microarray studies. Nucleic Acids Res, 2015. 43(7): p. e47.

22. Pino, L.K., et al., The Skyline ecosystem: Informatics for quantitative mass spectrometry proteomics. Mass Spectrom Rev, 2020. 39(3): p. 229–244.

23. Wagner, A., et al., Metabolic modeling of single Th17 cells reveals regulators of autoimmunity. Cell, 2021. 184(16): p. 4168–4185.e21.

24. Lu, X.M. and H. Long, Nicotinamide N-methyltransferase as a potential marker for cancer. Neoplasma, 2018. 65(5): p. 656–663.

25. Harbeck, N., et al., Urokinase-type plasminogen activator and its inhibitor type 1 predict disease outcome and therapy response in primary breast cancer. Clin Breast Cancer, 2004. 5(5): p. 348–52.

26. Buyuk, B., S. Jin, and K. Ye, Epithelial-to-Mesenchymal Transition Signaling Pathways Responsible for Breast Cancer Metastasis. Cell Mol Bioeng, 2022. 15(1): p. 1–13.

27. Sotiriou, C., et al., Gene expression profiling in breast cancer: understanding the molecular basis of histologic grade to improve prognosis. J Natl Cancer Inst, 2006. 98(4): p. 262–72.

28. Farmer, P., et al., Identification of molecular apocrine breast tumours by microarray analysis. Oncogene, 2005. 24(29): p. 4660–71.

29. van ’t Veer, L.J., et al., Gene expression profiling predicts clinical outcome of breast cancer. Nature, 2002. 415(6871): p. 530–6.

30. Müller, A., et al., Involvement of chemokine receptors in breast cancer metastasis. Nature, 2001. 410(6824): p. 50–6.

31. Coppé, J.P., et al., Senescence-associated secretory phenotypes reveal cell-nonautonomous functions of oncogenic RAS and the p53 tumor suppressor. PLoS Biol, 2008. 6(12): p. 2853–68.

32. Ibrahim, S.A., et al., Syndecan-1 is a novel molecular marker for triple negative inflammatory breast cancer and modulates the cancer stem cell phenotype via the IL-6/STAT3, Notch and EGFR signaling pathways. Mol Cancer, 2017. 16(1): p. 57.

33. Demaria, M., Senescent cells: New target for an old treatment? Mol Cell Oncol, 2017. 4(3): p. e1299666.

34. Piccinin, E., G. Villani, and A. Moschetta, Cholesterol-LXR axis in metabolic regulation of liver fibrosis and hepatocarcinogenesis. Trends Endocrinol Metab, 2026. 37(11): p. 1063–1077.

35. Kapoor, I. and U. Varshney, Diverse roles of nucleoside diphosphate kinase in genome stability and growth fitness. Curr Genet, 2020. 66(4): p. 671–682.

36. Finetti, F., et al., Prostaglandin E2 and Cancer: Insight into Tumor Progression and Immunity. Biology (Basel), 2020. 9(12).

37. Wang, S., et al., Pericyte NNMT drives an immune-hostile vascular niche and restricts cancer immunotherapy. Science Bulletin, 2026. 71(8): p. 1912–1916.

38. Sun, Y., et al., Treatment-induced damage to the tumor microenvironment promotes prostate cancer therapy resistance through WNT16B. Nat Med, 2012. 18(9): p. 1359–68.

39. Ganesan, R., et al., Taxane chemotherapy induces stromal injury that leads to breast cancer dormancy escape. PLoS Biol, 2023. 21(9): p. e3002275.

40. He, B. and R. Ganss, Pericyte phenotype switching in cancer. Trends Cancer, 2025. 11(9): p. 877–888.

41. He, B., et al., Selective tubulin-binding drugs induce pericyte phenotype switching and anti-cancer immunity. EMBO Mol Med, 2025. 17(5): p. 1071–1100.

42. Dalin, S., et al., Deoxycytidine Release from Pancreatic Stellate Cells Promotes Gemcitabine Resistance. Cancer Res, 2019. 79(22): p. 5723–5733.

43. Halbrook, C.J., et al., Macrophage-Released Pyrimidines Inhibit Gemcitabine Therapy in Pancreatic Cancer. Cell Metab, 2019. 29(6): p. 1390–1399.e6.

